# IntestLine: a Shiny-based application to map the rolled intestinal tissue onto a line

**DOI:** 10.1101/2022.10.26.513827

**Authors:** Altay Yuzeir, David Bejarano, Stephan Grein, Jan Hasenauer, Andreas Schlitzer, Jiangyan Yu

## Abstract

To allow the comprehensive histological analysis of the whole intestine in one image, the tissue is often rolled to a spiral before imaging. This Swiss-rolling technique facilitates robust experimental procedures, but it limits the possibilities to comprehend changes along the intestine. Here, we present IntestLine, a Shiny-based open-source application to map imaging data of intestinal tissues in spiral shape onto a line. The mapping of intestinal tissues improves the visualization of the whole intestine in both proximal-distal and serosa-luminal axis, and facilitates the observation of location-specific cell types and markers. In summary, IntestLine serves as a tool to visualize and characterize intestine in future imaging studies.

## Introduction

A comprehensive assessment of large organs is a key challenge in many biomedical research fields. Therefore, different strategies have been devised to reduce the spatial field of view for spatial analysis. A commonly used approach for the intestine research is the Swiss-rolling technique, which has been shown to facilitate the study of the intestinal structure along the proximal-distal axis in a single image^1^. Combination of the Swiss-rolling technique and multiplexed imaging approaches such as co-detection by indexing (CODEX) has allowed to identify intestinal immune and stromal cells and their location along the whole intestinal structure at single-cell resolution^2^. In addition, embedding the spiral shape of intestine on the 10x Visium slide has provided the spatial gene expression of all cells of the complete intestinal structure^3^. These cutting-edge techniques facilitate the identification of location-specific cell types, molecular markers and cell-cell interactions along the intestinal proximal-distal and serosal-luminal axis^3^. Yet, the visualization of the imaging data for the intestinal tissue as an artificial spiral (created by Swiss-rolling technique) is highly non-intuitive and limits the comprehension. Thus, a tool which maps rolling tissue to the natural linear coordinate system would be of great benefit for the research community.

A theoretical possibility to unroll the imaging data obtained from the Swiss-rolling technique, would be to fit the rolled tissue to the Archimedean spiral model (*r* = *a* + *b* * *θ*, where *r* is the distance from the center and *θ* is the angle from the horizontal axis, and *a* and *b* are constants), and to use the path length on the spiral as the position in a linear coordinate system. However, for the established experimental procedures, the distance between layers is often non-uniform along the coordinates of the spiral. Thus, a perfect spiral is an insufficient model for the rolled tissue.

A recent study by Parigi et al. ^3^ has reported a custom pipeline to convert the intestinal tissue in the spiral shape from 10x Visium technique into a linear coordinate system. The pipeline implements three steps: (1) Selection of the base layer using Photoshop (which will act as a skeleton for the reconstruction); (2) Ordering of base layer points from proximal to distal by constructing a shortest path from the defined start to the end; and (3) Assignment of all spots (cells) to the ordered base layer. The use of the ordering position of the based layer point as a (linear) coordinate allows the visualization of genes enriched in distal and proximal regions. Yet, while this pipeline is helpful, it is not available as an open-source tool. Furthermore, only about 100 points were used for the base layer, severely limiting the resolution. Therefore, new tools are demanded to visualize the whole intestine in a linear coordinate system with higher resolution.

Here, we present IntestLine, a Shiny-based tool to map intestinal tissue with spiral shape onto a line. We adopted the general strategy described in Parigi et al.^3^, which includes the manual selecting points for the base layer, the assignment of other spots (cells) to the closest proxy on the base layer and the visualization of the intestine in a linear coordinate system using the positioning implied by the coordinate on the base layer. To allow for a problem-specific resolution, IntestLine allows for the selection of a flexible number of points for base layer from the inner (distal) to outer (proximal) side of the image. The mapping can be exported for downstream process such as visualization of marker intensity or for analysis of other relevant parameters at single-cell resolution.

## Results

### Workflow of IntestLine

IntestLine is an open-source application and is implemented in a docker with the pre-installed Shiny app. To create linear visualization of the intestine from the images of a slice prepared by the Swiss-rolling method, users first upload a text file in the CSV format containing cell locations to the IntestLine application **(Figure 1)**. In order to assign all cells to the base layer, a center point needs to be selected by the user. Next the dataset will be uploaded into the Shiny app to allow users to select points for the base layer. IntestLine makes use of the order of points picked in the Shiny app to reconstruct the base layer of the linear coordinate system and thus it is important to select points from inner (distal) to outer (proximal) in order. The selected points for base layer could be downloaded as a text file and be re-uploaded for future analysis. After that, each spot (or cell) in the image will be assigned to the nearest base layer point, which has a larger distance to the center point than the query spot. Meanwhile the distance of the spot to the corresponding base layer point is calculated and will be later used as the thickness (as y-axis) in the linear coordinate system. Later, in order to remove noisy signals in the gap of two intestinal layers, for the group of spots assigned to the same base layer point, the Z-score of distances within the group of spots will be calculated. Users can define their own threshold on the thickness and the Z-score in the filtering step within the app. Finally, the rolled image will be converted into a linear coordinate system using thickness as y-axis, and the cumulative length of base layer points as x-axis. The linear mapping can be exported for further visualization and analysis.

**Figure 1.**
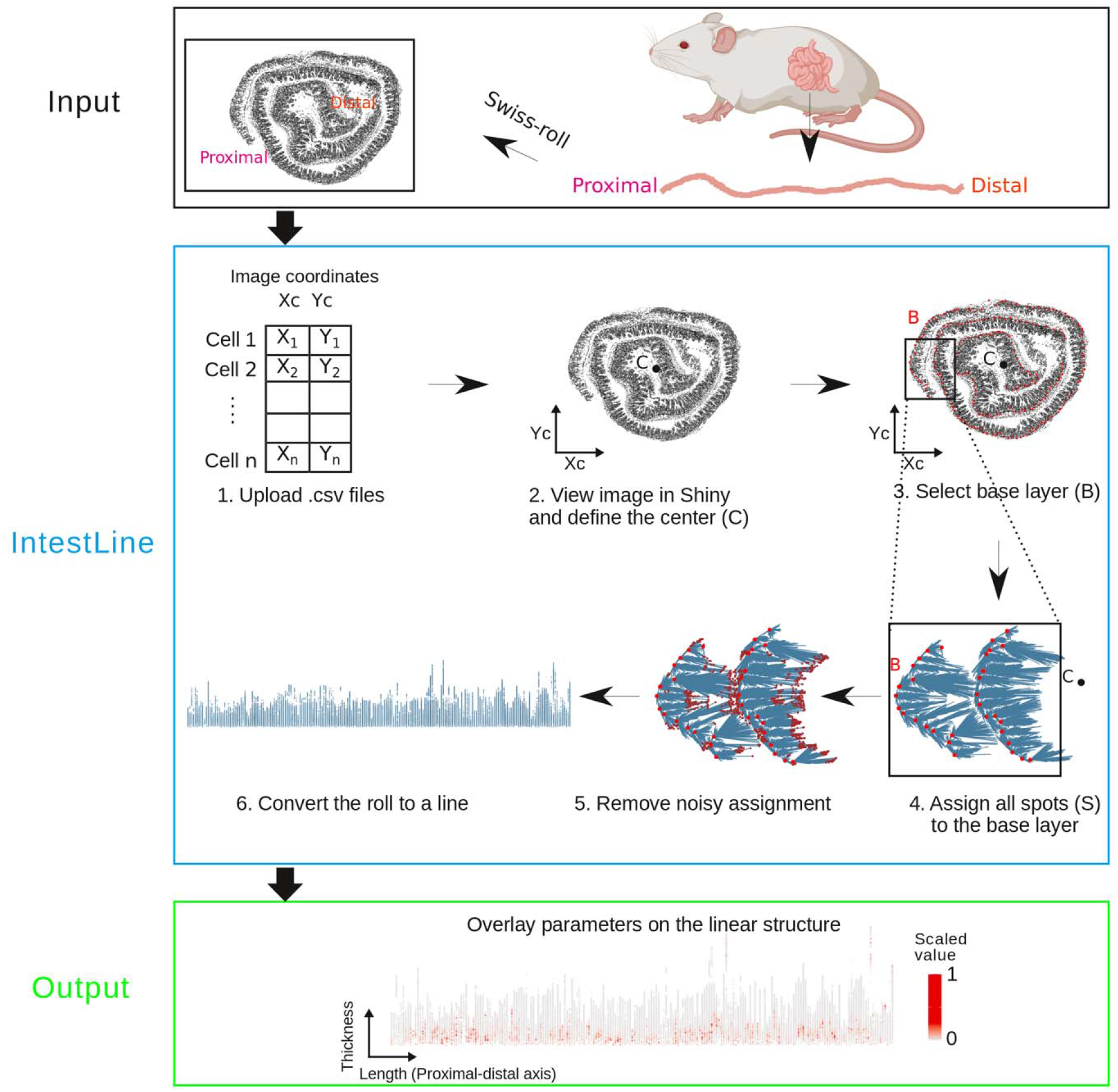
Workflow of IntestLine. After uploading the csv file containing xy-coordinates from individual cells, a center (C) point will be manually defined in the image. Next points to form a base layer (B, denoted in red in steps 3-5) from inner to outer will be manually selected in the implemented Shiny app. Later all cells or spots (S) in the image will be assigned to the nearest outer adjacent base layer point that is with larger distance to the center point. Finally, after setting the user-defined noisy threshold including distance to the base layer (thickness) and Z-scores (denoted in dark red in step 5). Here the Z-score of distance is calculated within a group of spots assigned to the same base layer point. The image will be automatically converted into a linear coordinate system with thickness as y-axis and cumulative length of base layer points as x-axis.

### Application of IntestLine

To evaluate IntestLine, we consider a CODEX image of the murine intestine that was prepared by Swiss-rolling technique and was stained with a 15-plex antibody panel. The resulting image was segmented using the CODEX processor V1.7, yielding a total of 150,793 cells (**Figure 2A**). First, we uploaded the file containing cell locations (xy-coordinates) exported from the CODEX processor into the IntestLine application. Next, we manually selected a base layer containing 1,059 points (**Figure 2B**). After assigning cells to the base layer (**Figure 2C**), we performed a stringent filtering by removing noisy assignment with thickness >1,000 or Z-score >2 (**Figure 2D-E**). As a result, more than 93% of cells were successfully assigned to the outer adjacent base layer (**Figure 2F**). Finally, we visualized the image in a linear coordinate system as shown in **Figure 2G**. The mapping of the intestine on a line allows us to observe clear thickness differences in proximal-distal axis. The results show that IntestLine is able to provide a high-resolution mapping of an image of a rolled intestinal tissue to a line.

**Figure 2.**
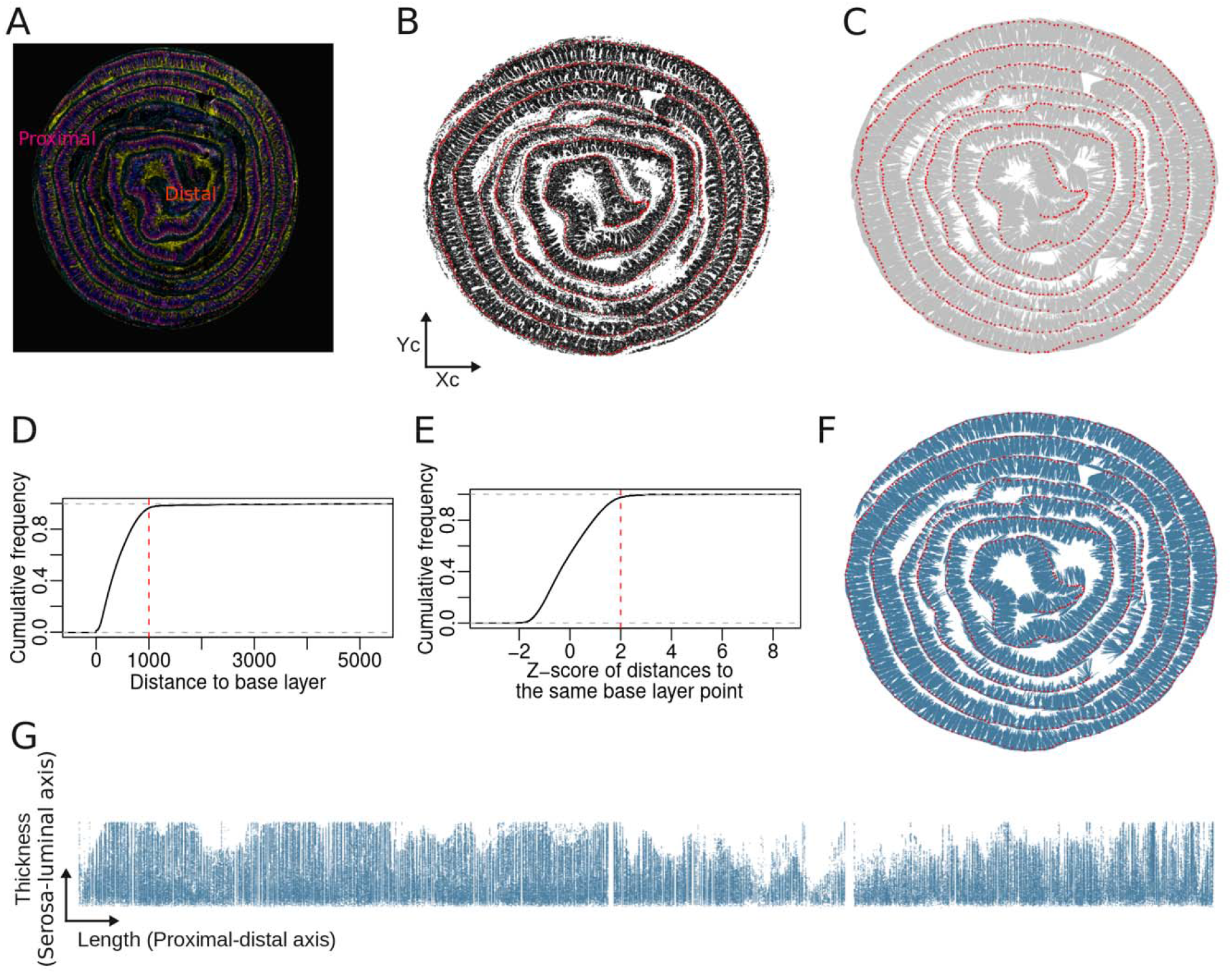
Pilot study of mapping an intestinal tissue onto a line. (A) The rolled intestinal tissue imaged by the CODEX technique. (B) Manually selected base layer (depicted in red) for the image. (C) Visualization of assigning spots to base layer before filtering process. The grey line connects the cell or spot to its corresponding base layer point (denoted in red). (D) Cumulative distribution of the thickness (distance of the spot to its corresponding base layer point). (E) Cumulative distribution of Z-score of distances within a group of spots assigned to the same base layer point. (F) Visualization of the successful assignment after filtering out assignment with thickness > 1,000 or Z-score > 2. (G) Converted linear coordinate system of the tissue. Thickness (y-axis) is the distance of the spot to its corresponding base layer point. Length (x-axis) is the cumulative length of base layer points.

### Capabilities of converted linear coordinate system to visualize fluorescent marker intensity

To demonstrate the importance of the data provided by IntestLine, we compared the fluorescent intensities of several markers between the original coordinate system and the converted linear coordinate system. While the direct visualization of the data does not facilitate the identification and quantification of trends, the unrolling provides a clear picture of the change along the serosal-luminal axis of the tissue (**Figure 3**). For example, Villin staining clearly highlighted the epithelium on top of the structure^4^, whereas olfactomedin-4 (Olfm4) and Ki67 staining accurately revealed the location of proliferating intestinal stem cells at the base of the crypts of the murine small intestine^5^. Meanwhile, lysozyme staining labeled Paneth cells at the base of distal intestinal region^6^. Moreover, the linear representation within the intestinal wall allowed the identification of regions with abnormal marker expression levels. For instance, the region highlighted in Figure 3B is with low expression of Ki67 and lysozyme, suggesting a disrupted structure in this neighborhood. In comparison, this information is less obvious from the intestinal image in the spiral shape. Taken together, the linear coordinate system would allow better visualization in both proximal-distal and serosa-luminal axes of the whole intestinal structure.

**Figure 3.**
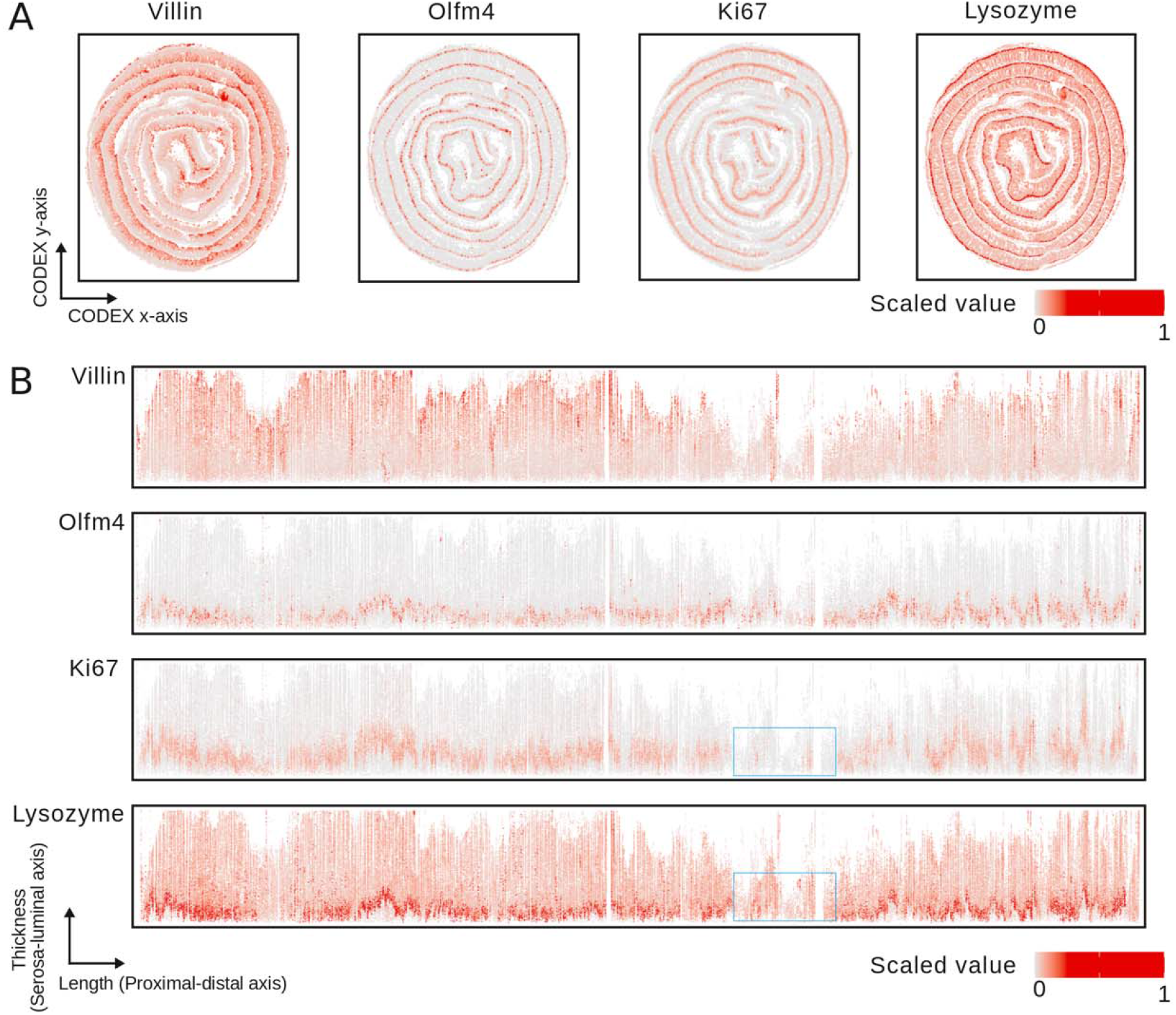
Visualization of fluorescent marker intensity on the tissue. (A) Fluorescent signal intensities of Villin, Olfm4 (intestinal stem cell marker), Ki67 (proliferation marker) and lysozyme in original CODEX xy-coordinates. (B) Fluorescent signal intensities of Villin, Olfm4, Ki67 and lysozyme in the converted linear coordinate system. The region lacking of Ki67 and lysozyme expression is highlighted in a blue box.

## Conclusion

The processing of imaging data is key for the assessment of biological processes. Here, we have presented IntestLine, one first open-source application to map rolled intestinal tissue images onto a line. We have shown that the mapping to a linear coordinate system facilitate the data visualization and the understanding of intestine anatomy as well as regions enriched with specific markers and cell types. Beyond this, the linear representations enable the embedding of all other available parameters generated by the CODEX processor or downstream analyses (e.g., cell clustering). Therefore, IntestLine application will provide a unique opportunity to characterize intestines in its natural linear shape for future mechanistic studies.

## Author contributions

J.Y. and A.S. conceived the project. D.B. provided CODEX images. J.Y. and A.Y. developed the pipeline. A.Y. designed the user interface. S.G. and J.H. provided critical feedback on the pipeline development. J.Y. and J.H. wrote the manuscript. All authors discussed the results and commented on the manuscript.

## Data availability statement

Source code can be found at Zenodo (https://doi.org/10.5281/zenodo.7081864) and Github (https://github.com/JiangyanYu/IntestLine). An open-source web application is available at FASTGenomics (https://beta.fastgenomics.org/a/intestline).

